# Pulsed laser lensing for phase modulation in electron microscopy

**DOI:** 10.1101/2025.06.12.659428

**Authors:** Daniel X. Du, Adam C. Bartnik, Cameron J. R. Duncan, Usama Choudhry, Tanya Tabachnik, Chaim Sallah, Yuki Ogawa, Ebrahim Najafi, Ding-Shyue Yang, Jared M. Maxson, Anthony W. P. Fitzpatrick

## Abstract

Phase contrast electron microscopy is fundamental for visualizing unstained biological specimens. Advances in electron detection have not yet overcome the low contrast caused by weak scattering. Here, we demonstrate that an orthogonal pulsed laser-electron beam interaction produces a pronounced peak phase shift of 430 radians through ponderomotive defocusing, leading to a maximum angular deflection of 45 µrad. Experiments encompassing a variety of probe pulse energies and pump positions verified the properties of the electron pulses in a range of pulse durations from 5.8 ± 1.9 ps to 13.4 ± 0.9 ps and a width of 15.0 ± 2.6 µm at the interaction region. The stability of the beam was also tested across 10 hours of cumulative acquisition time, with only small variations in laboratory conditions resulting in a gradually shifting baseline measurement. Pulsed laser lensing of the electron beam offers the potential for refinement in phase shift and electron beam shaping with careful consideration to the overlap between laser and electron pulses. Calculations of phase shifts across a wide experimental envelope show that poorly chosen laser parameters can generate large incoherent distributions at both 30 keV and 300 keV. Thus, a delicate balance between laser and electron widths and pulse durations must be struck to adequately achieve uniform phase shifts, particularly when singling out specific beamlets in the back-focal-plane.

## Introduction

Almost a century ago, magnetic lenses were first incorporated into an electron microscope(1). Since then, electron microscopy has advanced from barely surpassing the optical diffraction limit(2) to employing aberration corrected lenses that resolve features smaller than fifty picometers(3), close to the Bohr radius of a hydrogen atom. These extraordinary gains in precision, stability, and versatility underpin many imaging breakthroughs in materials science(3), nanotechnology(4), and structural biology(5).

Beyond static magnetic fields, electron beams can also be manipulated by time-varying electromagnetic fields, such as laser pulses, through the ponderomotive effect. Recent work shows that the optical field of a laser can be used to shape the beam(6) or impose precise phase shifts(7). The types of ponderomotive interactions can roughly be organized into two categories. The first is the Kapitza-Dirac effect, involving two-or-more beams traveling in different directions that impart discrete momentum shifts governed by multiples of the photon momenta(8, 9). The other is a simple beam-beam interaction that relies solely on electrons interacting with the field of the laser beams(6, 10–12). To avoid discrete momentum shifts, which may be deleterious in imaging applications(13), the second type of effect will be the focus of this publication. Physically, this translates to just using a single laser beam. The theoretical framework for this interaction is well established for many applications, from lensing to aberration correction(12, 14–18), and has been extensively simulated(19). Experimentally, the ponderomotive effect has been used for time-zero overlap calibrations(20–24) and, most recently, Mihaila *et al*. have demonstrated lensing and beam-shaping in a laser co-propagating and counter-propagating configuration with respect to the direction of electron travel(6). Deflecting an electron beam with light requires peak optical intensities on the order of GW/cm^2^(13). Such extreme intensities are most readily obtained with ultrafast laser pulses(10). For example, compressing just one microjoule of laser energy into a pulse lasting only a femtosecond generates an instantaneous power of one gigawatt.

Here, we conduct stroboscopic ultrafast experiments that explore how an orthogonally traveling pulsed laser beam, which is introduced into the microscope through standard optical windows, can induce pronounced spatial bending of electron packet trajectories (Fig. 1a). We extend this to a discussion on the impact of overlap between the electron and laser beam. The pulsed laser lensing corresponds directly to phase shifts in the electron wavefront. This proof-of-concept experiment shows that laser parameters set outside the microscope can readily induce a ponderomotive potential and produce phase shifts of several hundred radians in the electron beam. We also include a discussion on the necessities behind transferring this experiment from the scanning electron microscope (SEM) to the transmission electron microscope (TEM), which is known to have challenges in matching the electron beam currents seen in normal field emission gun (FEG) and thermionic emission modes.

**Fig. 1.**
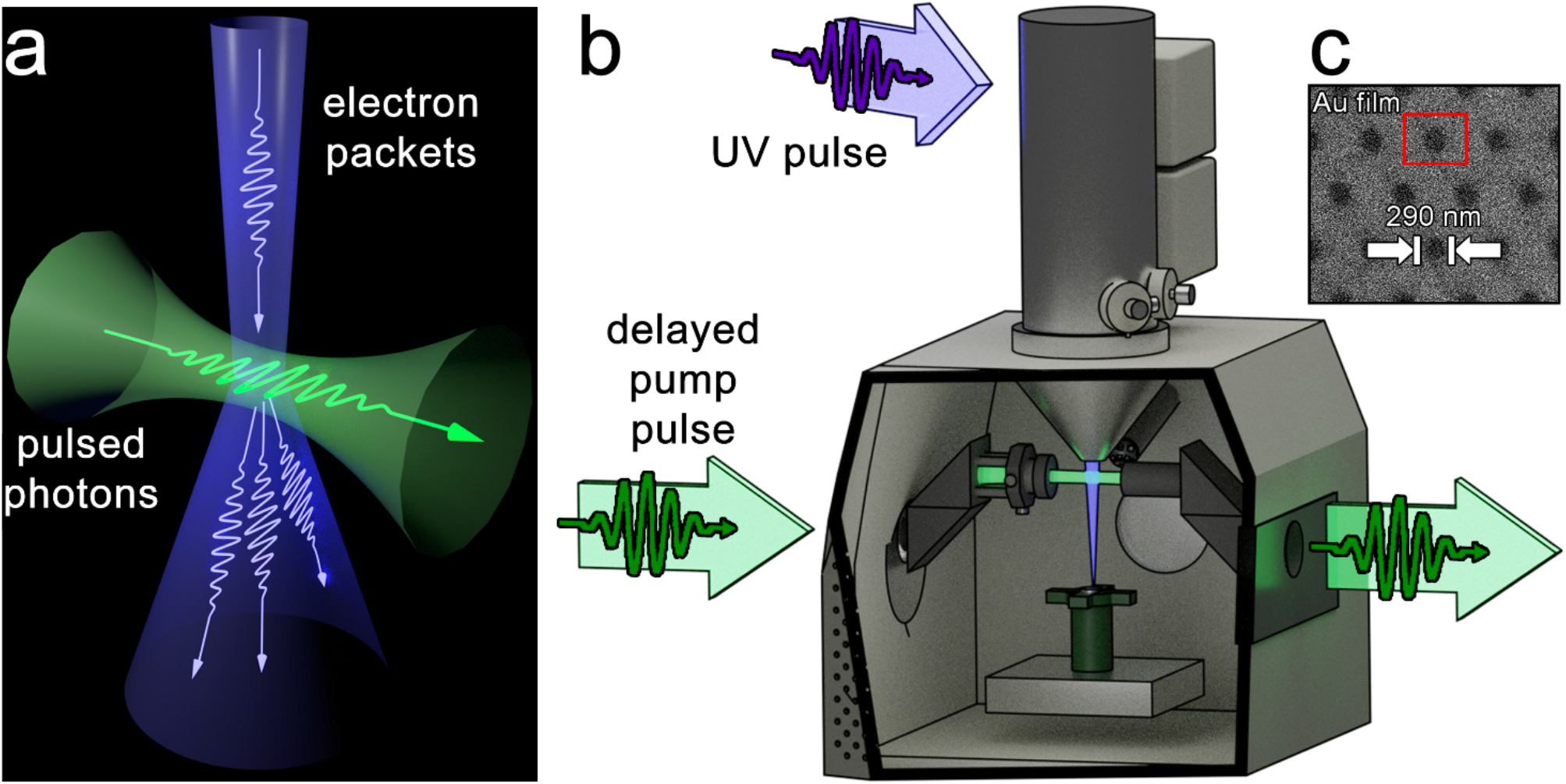
Schematic of the laser optics integrated into a USEM (ultrafast SEM) to allow for ponderomotive experiments. **a**, A pulsed laser (green) intersects with a strobed electron beam (purple) orthogonally, making each interacting electron packet behave as though it passes through a concave cylindrical lens. The ponderomotive effect of the pulsed laser defocuses the electron beam and simultaneously imposes a phase shift. **b**, A cross-section of the USEM that shows the optical connections. The electron beam is typically focused to a tight point to achieve high resolutions, but the interaction expands a subset of the focused electrons. A set of periscopes were installed to allow for the longest propagation distance post-interaction but are not necessary for the ponderomotive interaction itself. **c**, An image of the gold foil (290 nm diameter holes) used for experiments, with the region of interest highlighted by a red box.

## Results

To quantify the momentum kick generated by the ponderomotive effect, we back-calculate the phase shift using a set of equations(6) for the case of orthogonal beams, where the laser takes the form:

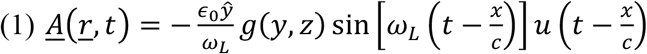

Here, *c*, and *ϵ*_0_ are universal constants, *g*(*y, z*) is the TEM_00_ gaussian profile, and *ω*_*L*_ is the frequency of the light. We derive the momentum shift from 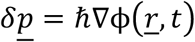 (see Supplementary Note 2) using a path integral, which is immediately applied to classically treated electrons with a known movement vector. A varied set of input movement vectors results in a distribution of the electron beam at the imaging plane. To mimic the experiment and the nature of the detector, a spatial filter is required to detect the change in the electron beam.

For this first experiment, we measure the ponderomotive defocusing of the electron beam by imaging 290 nm holes in a gold foil with an Everhart-Thornley Detector and observing the resulting beam spread. Secondary electrons are generated at the surface of the specimen by the focused electron beam and drawn into an Everhart-Thornley detector (ETD) positioned towards the back of the specimen chamber (Fig. 1a). We purposefully choose the ETD over the silicon-diode scanning-transmission electron microscopy detector as a spectral filter placed after the ETD phosphor is more effective in suppressing signal from scattered laser light. Images are formed by rastering the beam across the specimen on a pixel-by-pixel basis. The induced phase shifts are sufficiently large that the corresponding momentum changes are detectable, as the ponderomotive effect pushes the electron away from the region of highest photon density. To demonstrate elastic interactions between a pulsed laser beam and strobed electron packets, we generate photoelectrons (electron energy 30 kV, pulse duration of 6.8 ± 0.9 ps) with femtosecond ultraviolet (UV) pulses (0.5 nJ/pulse, 212 fs fwhm, 1 MHz repetition rate) from a zirconium-coated tungsten tip and intersect them orthogonally with green laser pulses (*λ* =512 nm). The interaction volume is approximately 14 × 14 × 14 μm^3^. The built-in picoammeter and Faraday cup were incapable of providing a read-out of the beam current at the UV pulse energy values used in this publication, though we can confirm that the ETD was required to be at 100% gain to resolve the specimen.

Given that the intensity changes induced in the electron beam by the ponderomotive effect are quite small, we subtract micrographs taken without electron-laser interaction from those recorded at time zero; the exact moment when electron packets and pulsed photons interact directly. The images are median-filtered to suppress any anomalous high-frequency features that may arise due to noise.

Difference images of single holes (Fig. 2a, *left*) are compared with simulated difference images (Fig. 2a, *right* and Supplementary Note 1). The ponderomotive effect deflects the electrons along what appears to be an axis 41° from the normal to the laser beam’s propagation direction (Fig. 2a, *left*), primarily due to severe astigmatism in the beam due to misalignments discovered afterwards (Supplementary Note 2). Counts integrated along this line are plotted (Fig. 2b). The smaller lobe to the bottom left has diminished intensity and the electrons are pushed into the top right corner, with a peak mean difference intensity of 1.5%. The simulated image provides a similar shape profile (Fig. 2a, *right*), but the predicted signal is somewhat muted, suggesting the effect is stronger than expected (Fig. 2b).

**Fig. 2.**
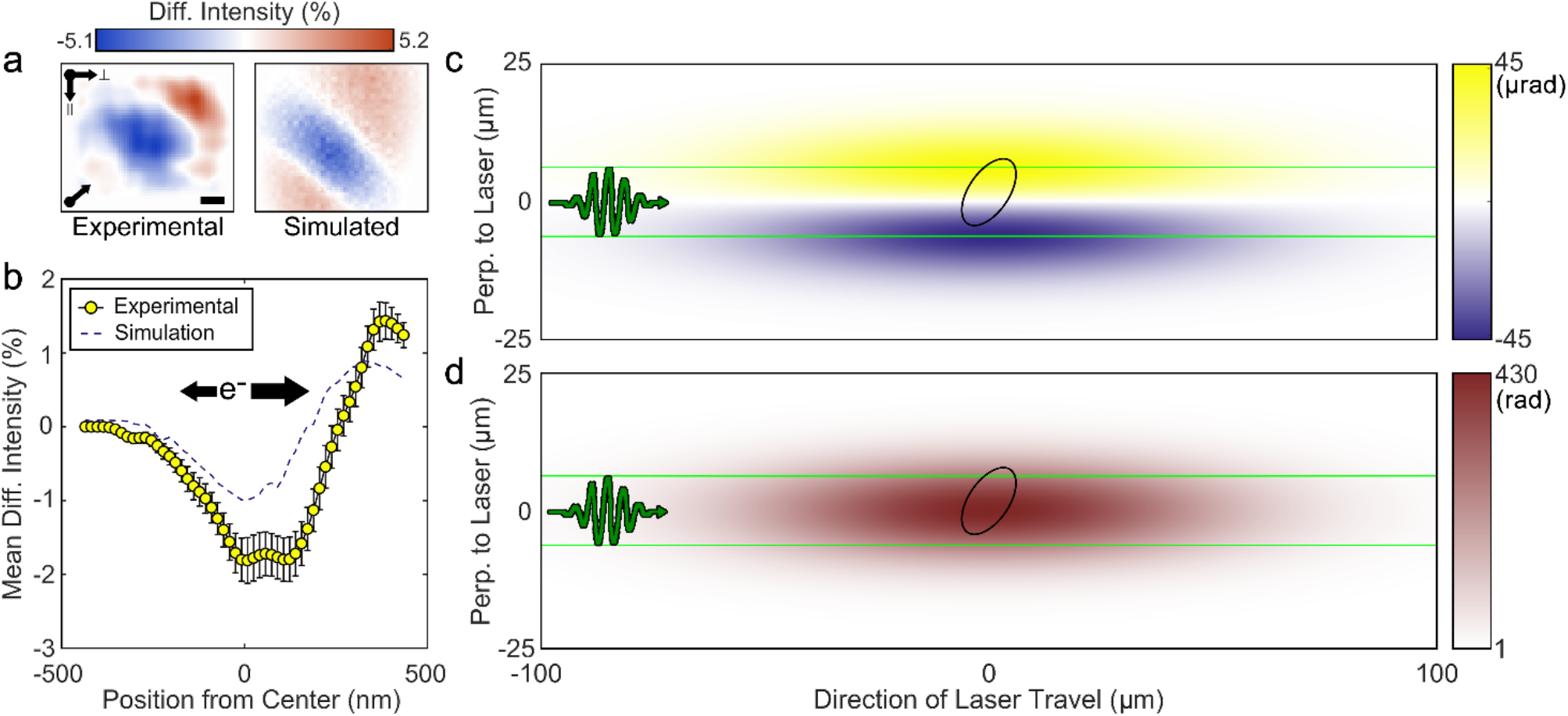
Pulsed-laser lensing of an electron beam using the ponderomotive effect. **a**, Difference images were generated by subtracting the negative-time reference image, acquired 21-67 ps before the photons arrive, from the time-zero image, when the photon pulses overlaps the electron packets. After noise filtering, these experimental difference images (b, top) are compared with simulated counterparts (b, bottom). The colorbar indicates the difference intensity in percentage of peak image counts. Axes are provided to indicate directions parallel and perpendicular to the laser Poynting vector. A third arrow is used to show the near-diagonal direction of laser lensing due to the ponderomotive effect. Scale bar, 100 nm. **c**, A line profile of the difference intensity along the axis of pulsed-laser lensing due to the ponderomotive effect. The direction along which electrons are preferentially pushed is indicated by the larger black arrow. Experimental data is shown as filled yellow circles (±S.E.M.) with simulated data shown as a dashed line. **c**, Momentum shift (µrad) produced by the ponderomotive effect in the axis orthogonal to both beams. A 500-fs laser pulse (green wave packet, propagating left-to-right) intersects an electron packet that moves into the page (black outline). The continuous green lines mark the 12.5 µm laser-beam diameter, while the black ellipse (major axis ∼15 µm) shows the simulated electron-beam profile calculated from the experimental parameters in Supplementary Table S1. **d**, The ponderomotive effect generated by the pulsed laser (green wave packet propagating left-to-right; 12.5 µm beam bounded by solid green lines) imparts an equivalent phase shift of roughly 430 radians to the electron packets traveling into the page (black outline).

The unexpected astigmated nature of the electron beam resulted in an optimization in maximizing the number of momentum-shifted electrons that could be detected, yielding a difference that is easier to quantify. For each experiment, images are acquired such that the ratio of the noise and the contrast between the maximum and minimum intensity values were limited to 0.01, or below. From the total compiled images, a final image is obtained through a least-squares difference (Fig. 2a, *left*). By correlating empirically found electron beam properties (Supplementary Note 1) to simulated data, we expect to understand the phase profile imparted on the electrons passing through the laser beam. Here, we have determined the matching simulated parameters by the magnitude and angle of the electron deflection. The parameters that most closely matched the acquired image were then checked against the experimental parameters determined previously and found to be within the expected ranges (Supplementary Note 1). Using the characterized electron beam parameters (Supplementary Table S1), we observe the expected cylindrical-like concave lens behavior, with a focal length of approximately −42 ± 14 mm.

The momentum kick imparted by the pulsed ponderomotive potential to the electron beam (Fig. 2c and Supplementary Note 2) is also calculated from the same simulations. The magnitude of the resulting momentum shift, approximately 90 µrad, is strongly anisotropic and lies mainly along the major axis of the black ellipse bounding the electron beam in Fig. 2c. This directional bias explains the weighted lobe pattern seen in Fig. 2a. Converting the measured momentum shift to its equivalent phase reveals that the ponderomotive interaction imposes a peak phase of roughly 430 radians on the electron beam (Fig. 2d), which far exceeds the required π/2 radians for phase interference, but is necessary to visualize on the ETD.

To probe how the ponderomotive effect varies with electron-pulse length, we tuned the pulse duration by changing the UV excitation energy. Lower UV energies produce shorter electron packets that carry fewer electrons, which lessens space-charge repulsion and improves beam coherence(25). In the case of the ETD, the minimum UV pulse energy that generated an observable and resolvable electron beam was at 0.25 nJ/pulse. Scattered laser light introduces a significant amount of noise, even after spectral filtering. This signal-noise ratio varies from day-to-day depending on the health of the tungsten-ZrO tip, resulting in the aforementioned 1% noise acquisition requirement. At a UV pulse energy of 0.25 nJ, the electrons form packets only 5.8 ± 1.9 ps long and exhibit the largest intensity change, −4.1 ± 1.1% (Fig. 3a, *top*). Increasing the UV pulse energy to 0.5 nJ stretches the pulse to 7.1 ± 1.1 ps and lowers the intensity change to −3.3 ± 0.2%, averaged over two measurements (Fig. 3a, *middle*). Further increasing the pulse energy to 1.0 nJ lengthens the electron packet to 13.4 ± 0.9 ps, with the intensity change falling to −3.0 ± 0.1% (Fig. 3a, *bottom*). Thus, raising the UV energy from 0.25 nJ to 1.0 nJ more than doubles the electron-pulse duration while the magnitude of the intensity difference steadily decreases. We attribute the inverse relationship between intensity and UV pulse energy to an increase in overlap between the electron beam and the laser beam at lower electron beam durations. The impact of overlap is further discussed in a later section.

**Fig. 3.**
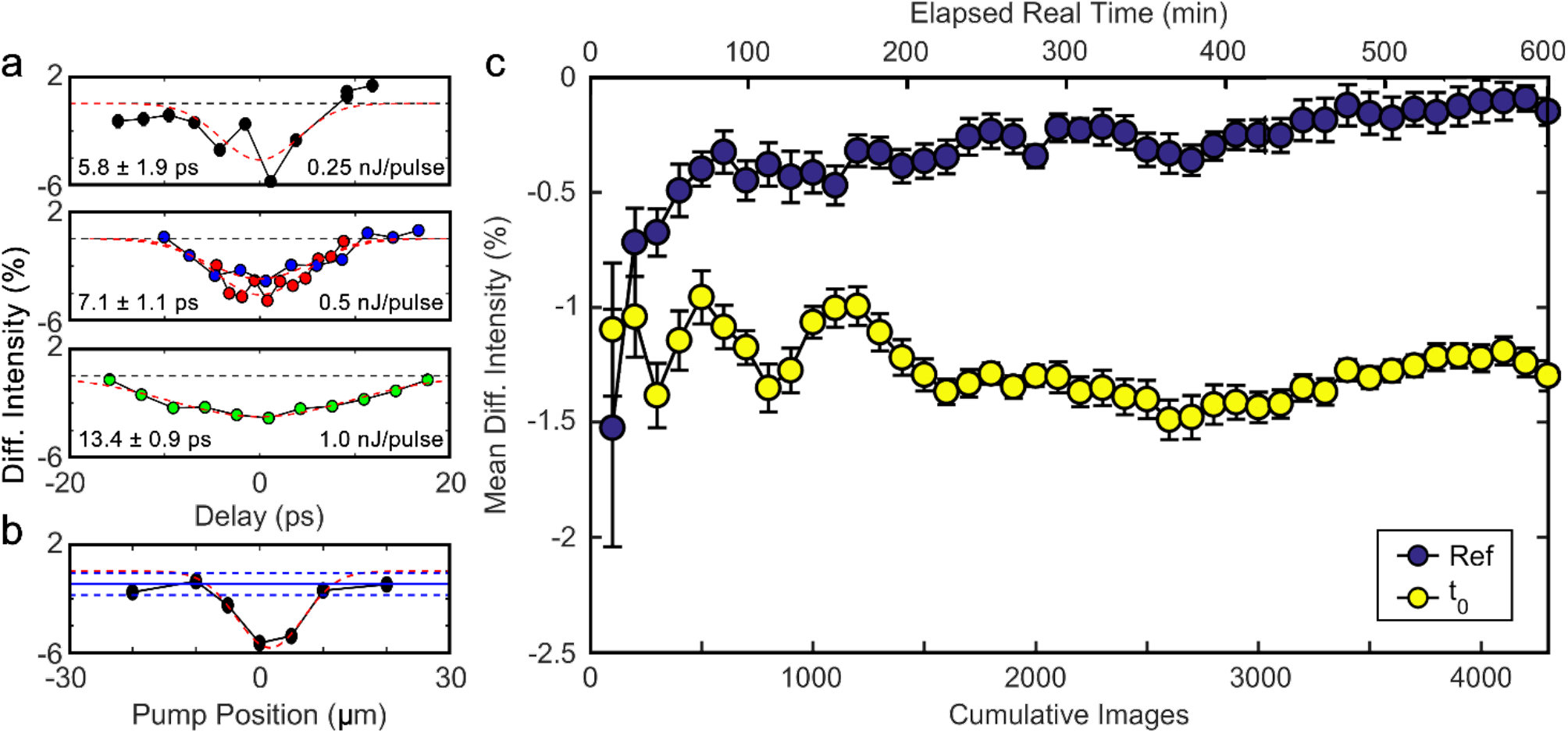
Maximizing pulsed-laser lensing by exploiting the ponderomotive effect over long periods of time. **a**, Increasing the UV pulse energy (*top*, 0.25 nJ/pulse → *middle*, 0.5 nJ/pulse → *bottom*, 1.0 nJ/pulse) lengthens the photoelectron packet (*top*, 5.8 ± 1.9 ps → *middle*, 7.1 ± 1.1 ps → *bottom*, 13.4 ± 0.9 ps). An approximate twofold increase in electron pulse duration weakens the ponderomotive effect, as evidenced by a steady reduction in the intensity difference (*top*, −4.1 ± 1.1% → *middle*, −3.3 ± 0.2% → *bottom*, −3.0 ± 0.1%). **b**, A strong ponderomotive effect requires the pulsed laser and the electron packets to have maximum spatial overlap. Shifting the laser beam perpendicular to its axis illustrates how even slight misalignment weakens the interaction. The solid blue line shows the mean signal in the negative-time reference images, while the flanking dashed blue lines denote its error bounds within a standard error to the mean (± S.E.M.). **c**, Demonstration of stability of pulsed-laser lensing due to the ponderomotive effect. Filled yellow circles (± S.E.M.) denote the mean difference intensity at time zero, while filled blue circles (± S.E.M.) indicate the mean difference intensity at −21 ps.

Beyond electron-pulse duration, spatial precision proves equally critical for maximizing the ponderomotive effect. The intensity modulation drops precipitously outside the narrow ∼20 µm interaction region where the electron and laser beams intersect (Fig. 3b), demonstrating that even minor alignment errors dramatically suppress the interaction strength. This spatial sensitivity, combined with the pulse duration dependence, establishes that optimal ponderomotive coupling requires both ultrashort electron pulses and precise beam overlap.

In a second experiment focused on the stability of the effect, we acquired 4,200 consecutive difference images (Fig. 3c). The integration time for each image was 2.8 seconds at a UV pulse energy of 0.75 nJ/pulse across three timepoints, resulting in a total integration time of 10 hours. The resulting contrast difference is a direct result of a redistribution of electrons from the ponderomotive effect. As previously discussed, any observable momentum shift suggests a strong phase shift. Over the 10-hour measurement period, the contrast of the effect varies by less than 10% from its mean value, which is −1.27% ± 0.13% (Fig. 3c, yellow circles). Difference images recorded when the laser and electron beam do not interact (*i*.*e*. when the retroreflector is moved far away) converge to a mean difference intensity of zero, as expected (Fig. 3c, blue circles). This data demonstrates that the effect is relatively stable for long periods of time, provided that no external factors result in a spatial misalignment of either beam. There are still sources of instability, as is shown in the data itself in the form of the slowly moving reference, which we have attributed to a slowly weakening electron beam, resulting in a marginally higher noise floor by the end of the scan. Similarly, the drift in the t_0_ data seems to suggest some small movement in the laser alignment. We have found that the laser does not move more than 5 microns over the period of a week unless the airbags inflate. Critically, these sources of instability may play small roles in the momentum shift but would result in significant shifts in phase interference due to the cyclical nature of phase. Many of these instabilities can be addressed with the installation of vibration-dampening systems and stronger environmental controls, but should be carefully considered, nonetheless. Additional controls and time-zero determinations are further explored in Supplementary Note 3.

## Discussion

We have previously mentioned that overlap between the electron beam and laser beam plays a significant role in the distribution of phase shifts imparted during the interaction. In the case of a phase plate, the distribution of the phase shift should be as narrow as possible, otherwise contrast enhancement is diminished, at best, or obfuscated, at worst. The easiest method of shortening the electron beam pulse duration is to reduce the number of electrons *per* packet and eliminate coulombic repulsion, though travel through various optics may result in an elongated electron pulse. A naïve linear fit to the pulse duration in Figure 3a suggests an instrument response of 2.4 ± 1.1 ps. While this is a poor response for the purposes of increased interaction, we attribute this increased response to poor electron lens settings and vibrations given that other ultrafast electron microscope (UEM) systems have shown laser-limited instrument responses(26, 27). As a thought experiment, we have fixed an example electron beam at 1 picosecond duration and a width of 10 microns. From here, the variable parameters are the electron energy, laser diameter, duration, and pulse energy.

Figure 4a shows simulated (with the same formulae as previously mentioned) pulse energies required to reach a mean phase shift of π/2 with the variation in laser parameters. These energies predictably increase as laser duration and laser diameter increase in size. Importantly, the differences between 30 keV and 300 keV electrons are also shown at the colorbar, with the 300 keV electrons requiring more pulse energy to reach a mean phase shift of π/2. The near-linear relationship between laser energy density and electron beam density is also represented by the nearly overlapping contour lines. However, the differences between 30 keV and 300 keV electrons are more apparent in Figure 4b, which represents the distribution of the phase shift at various laser properties. As can be clearly seen, a more spatiotemporally compressed laser pulse results in a significant increase in the phase distribution, to the point that the root-mean-square (r.m.s.) deviation from the mean exceeds the average phase shift. However, the change from 30 keV to 300 keV does not affect the r.m.s. deviation for a π/2 mean to a significant degree at short laser pulse durations. Furthermore, this discrepancy is diminished at large laser pulse diameters, which matches the physical expectation that the energy density largely controls the phase distribution. A broad laser peak would therefore appear as a minimally varying field, regardless of electron energy.

**Fig. 4.**
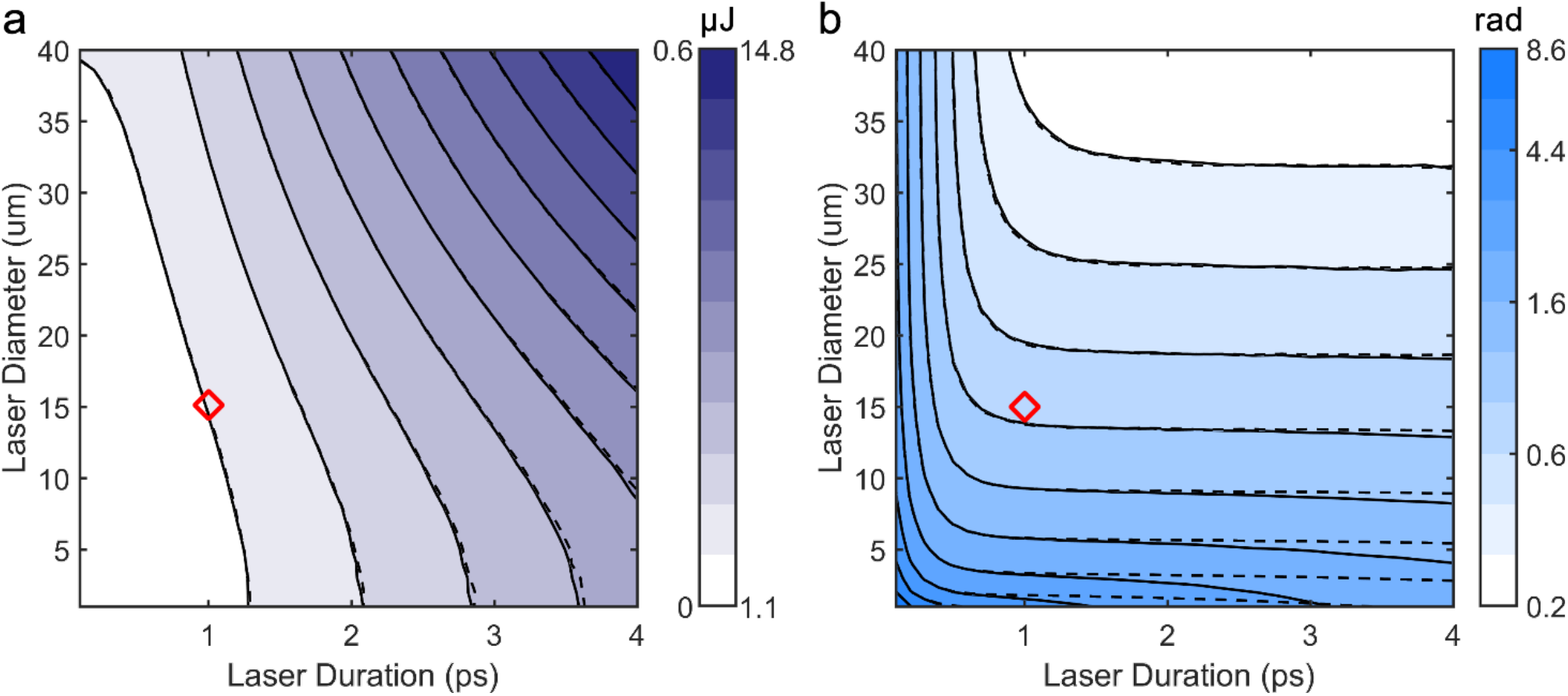
Simulated properties of an electron beam post-laser interaction at 30 keV (solid) and 300 keV (dashed). An electron pulse duration of 1 ps and a central unscattered beam diameter of 10 µm was assumed for all simulated values in this figure. Here, the laser parameters are varied to reach a mean phase shift of π/2. **a**, The required pulse energy in microjoules. The left (right) of the colorbar corresponds to 30 keV (300 keV) electrons and is on a linear scale. **b**, The r.m.s deviation from the mean phase shift. The colorbar values are displayed in radians but positioned on a log_10_-scale.

From the simulated plots, we can pick an example 15 micron laser diameter and a minimum of 1 picosecond pulse duration (marked with a diamond), leading to a r.m.s. deviation of around 0.6 radians. Pulse-stretching gratings with large group dispersion delays are effective enough to create short electron pulses and long laser pulses for these future experiments. Despite the classical particle analysis, this simulation relies upon one constraint: a coherent wavefront for each individual electron. However, we do not expect this to differ from any other requirement where weak scattering differences are observed from similar atomic elements (*i*.*e*. the case for biological specimens), which by necessity require a coherent electron beam. The primary form of instability would be in the alignment of the laser beam, which can be carefully controlled within a less vibration-prone environment, though the beam has been shown to be relatively stable (Fig. 3c). It does however require a pulsed electron beam in all imaging modalities and therefore imposes a more difficult-to-address constraint.

The overarching concern with UEM is undeniably beam current. As we have previously established, there is a strong inverse relationship between the pulse duration and the number of electrons *per* pulse generated at the cathode due to coulombic repulsion. This coulombic repulsion also plays a deleterious role in resolution. There is thus a conflict of interest between the beam current and resolution. The only method of solving this is to maintain single-electron or low-electron pulses and increase the repetition rate (or duty cycle) of the laser. In conventional UEM, these repetition rates are limited to kHz and low MHz frequencies to accommodate the relaxation time of photoexcited specimens(28–30). Even then, higher resolutions have already been demonstrated with 1.1 nm at 200 kHz and 10s acquisitions in the ultrafast TEM and 10 nm at 25.2 MHz in the ultrafast SEM(31–33), all of which have been direct acquisitions and do not involve any averaging techniques as in the cases of single-particle analysis or tomography. Where there is no such requirement for specimen photoexcitation and relaxation, duty cycles can be sharply increased to GHz frequencies. This may also confer the added benefit of diminishing the rate of beam damage as the electron emission becomes more regimented rather than stochastic(34, 35), though recent papers have suggested that such improvements are unlikely(36, 37).

In conclusion, we demonstrate that a femtosecond laser, introduced orthogonally to a pulsed electron beam, functions as a tunable ponderomotive lens(11). This free-space interaction imposes phase shifts of roughly 400 rad, greatly in excess of the π/2 threshold for strong interference, and imparts a stable angular kick of approximately 90 µrad that is stable for many hours. The lensing strength scales with electron-pulse duration and beam overlap, establishing clear experimental guidelines for optimum coupling in a transmission electron microscope (10, 38). With all parameters adjustable outside the electron microscope, this approach introduces a flexible, contact-free laser lensing technique that augments conventional electron optics. External optics allow the pulsed-laser phase plate to precisely shift its focus, keeping the laser-electron interaction aligned with the moving back-focal-plane crossover during sample tilting in pulsed-beam cryo-electron tomography (39), thus preserving optimal phase contrast and image quality. In addition to enhancing phase contrast in cryo-electron tomography (7), tailored laser fields could correct residual aberrations in real time (16, 17), narrow the energy spread for improved monochromaticity (40), or provide adaptive beam shaping for ptychography (41) or relativistic electron optics (42). Laser-based lensing thus provides a versatile approach for all-optical aberration correction and dynamic beam conditioning, enhancing and expanding the capabilities of traditional electron microscopes.

## Materials and Methods

The experiment consists of a pulsed laser and scanning electron microscope. The laser is an NKT Photonics Aeropulse FS20-050 operating a fundamental wavelength of 1030 nm and frequency doubled with a second-harmonic-generation module to 515 nm. The output power is 9W. The repetition rate of the experiment is set to 1 MHz for optimum power-per-pulse. The beam is initially passed through a 90:10 beam splitter. The weaker leg (probe) is further frequency doubled to 257 nm and modulated with waveplates and polarizers to generate photoelectrons at the ZrO coated tungsten FEG tip cathode *via* the photoelectric effect(43). To synchronize the laser pulse with the electron pulses, the stronger leg (pump) is delayed with a retroreflector and motion stage (mks IDL225-600LM). Due to spurious losses on the table from reflections and transmission, only 3.2 W of pump light enters the SEM. Once within the SEM, the pump is focused into the electron-photon interaction zone with a waist diameter of 12.5 µm (see Supplementary Note 2).

The microscope is a TFS Nova NanoSEM and, in photoelectron mode, operates at 30 keV accelerating voltage, 0.1 A tip current, and 1230 V in the C1 lens. A 1000 µm aperture is used to optimize the beam current, simultaneously setting the electron beam size to 12-24 microns at the interaction zone. The airbags were disabled to ensure laser positional stability, with less than 5 microns of drift after 7 days of experimentation. A standard silicon photoexcitation experiment was performed to identify the temporal overlap between electron and laser pulses on the delay stage (see Supplementary Note 2)(33). To ensure no side-lobes were produced for each electron pulse, the optical port originally at the cathode was replaced with a UV anti-reflective-coated window (TSL VPZ16QBBAR). To prevent the pump light from saturating the Everhart-Thornley Detector, a spectral filter (Asahi Spectra XVS0490 shortpass 490 nm) was fixed behind the light-pipe.

The holey gold spatial filter (Quantafoil HexAuFoil) with 290 nm holes is conductively bound to a degenerate p-type B-doped silicon (MTI Corporation, < 5 Å roughness, 0.001-0.005 Ω·cm resistivity) platform serving as a smooth conductive dark background and settled 10 mm/7mm below the polepiece/ponderomotive interaction zone. Images were collected at 10kx to 20kx magnification and in a narrowed ROI encapsulating one unit cell of the repeating holey pattern. Each acquisition was subtracted against a reference timepoint after drift correction. Difference images were achieved by minimizing the summed r.m.s. deviation. A second reference timepoint was acquired to serve as a control. This establishes the differences at time-zero are statistically different from any noise in the reference images, as is clearly shown in Figure 3c. A second control was performed by physically moving the specimen such that, if the effect was related to some surface phenomena, time-zero would also move. No such shift in time-zero was observed and the result of this experiment is shown in Supplementary Note 3.

## Acknowledgements

This work was supported by the Chan Zuckerberg Initiative, Visual Proteomics Grant under award no. 2021-234816.

## Notes

**Author contributions** D.X.D., A.C.B., C.J.R.D., E.N., D.Y., J.M.M., and A.W.P.F. conceived the experiments. D.X.D., T.T., C.S., and A.W.P.F. designed and fabricated components. D.X.D. built the optical set-up and carried out the experiments and data analysis, supported by A.C.B., U.C., E.N., D.Y., J.M.M., and A.W.P.F. Y.O. helped prepare figures. The manuscript was written by D.X.D., A.C.B., J.M.M., and A.W.P.F. after discussions with, and input from, all authors. A.W.P.F. supervised the project.

### Competing Interest Statement

The authors have declared no competing interest.

### Summary of Updates

This version of the manuscript has been revised to include additional data and a lengthier discussion.

