## Supplementary Information for "Pulsed laser lensing for phase modulation in electron microscopy"

Daniel X. Du[1,2,3], Adam C. Bartnik[4], Cameron J. R. Duncan[5], Usama Choudhry[6], Tanya Tabachnik[1], Chaim Sallah[1], Ebrahim Najafi[7], Ding-Shyue Yang[8], Jared M. Maxson[4*] & Anthony W. P. Fitzpatrick[1,2,3*]

[1] Mortimer B. Zuckerman Mind Brain Behavior Institute, Columbia University, New York, NY 10027, USA.

[2] Department of Biochemistry and Molecular Biophysics, Columbia University, New York, NY 10032, USA.

[3] Taub Institute for Research on Alzheimer's Disease and the Aging Brain, Columbia University Irving Medical Center, 630 West 168th Street, New York, NY 10032, USA.

[4] Cornell Laboratory for Accelerator-Based Sciences and Education, Cornell University, Ithaca, New York, NY 14853, USA

[5] Università degli Studi di Milano-Bicocca, Milan, Italy

[6] Department of Mechanical Engineering, University of California, Santa Barbara, CA 93106, USA

[7] The Chemours Company, Wilmington, DE 19899, USA

[8] Department of Chemistry, University of Houston, Houston, TX 77204, USA

*

### Supplementary Note 1. Simulations

#### 1.1 Derivation

The phase shift (as provided by an integral of the Hamiltonian) is:

$$(1)\ \phi(\underline{r}) = -\int dt' \frac{1}{\hbar} U(\underline{r}(t'), t')$$

where $U$ is the potential the electron passes through. The interaction potential for the ponderomotive effect is[1]:

$$(2)\ U(\underline{r}, t) = \frac{e}{2m_e\gamma}\left[A_x^2(\underline{r}(t), t) + A_y^2(\underline{r}(t), t) + \frac{1}{\gamma^2}A_z^2(\underline{r}(t), t)\right] + e\underline{v}\cdot\underline{A}(\underline{r}(t), t) - e\Phi(\underline{r}(t), t)$$

where $e$ and $m_e$ are the constants for the charge and mass of an electron respectively. $\gamma$ and $\underline{v}$ are the Lorentz factor and the velocity of the electron as dictated by the acceleration voltage. $\underline{A}$ and $\Phi$ are the vector and scalar potentials of the laser. We adopt the gauge where the scalar potential is zero and the high frequency photon oscillation negates the dot product term (besides being orthogonal). The vector potential of the laser beam is:

$$(3)\ \underline{A}(\underline{r}, t) \approx -\frac{\epsilon_0\hat{y}}{\omega_L} g(y,z)\sin\left[\omega_L\left(t - \frac{x}{c}\right)\right] u\left(t - \frac{x}{c}\right)$$

with $\epsilon_0$ as the field strength coefficient, $c$ as the speed of light, $\omega_L$ as the frequency of the photon, $g$ as the spatial distribution perpendicular to the travel direction, and $u$ as the temporal distribution. Thus, the phase is:

$$\begin{aligned}(4)\ \phi(\underline{r}, t) &\approx -\frac{1}{\hbar}\frac{\epsilon_0^2}{\omega_L^2}\frac{e^2}{2m_e\gamma}\int dt' g^2(x,y,z)u^2\left(t' - \frac{x}{c}\right)\sin^2\left[\omega_L\left(t' - \frac{x}{c}\right)\right] \\ &= -\frac{1}{\hbar}\frac{\epsilon_0^2}{\omega_L^2}\frac{e^2}{2m_e\gamma}\int dt' g^2(x,y,z)u^2\left(t' - \frac{x}{c}\right)\left\{1 - \cos\left[2\omega_L\left(t' - \frac{x}{c}\right)\right]\right\}\frac{1}{2} \\ &\approx -\frac{1}{\hbar}\frac{\epsilon_0^2}{\omega_L^2}\frac{e^2}{4m_e\gamma}\int dt' g^2(x,y,z)u^2\left(t' - \frac{x}{c}\right)\end{aligned}$$

We can assume the oscillatory sine and cosine terms do not contribute due to the high frequency oscillations of the photons. The laser pulse energy when integrated to infinity is given by:

$$\begin{aligned}(5)\ E_L &= c\varepsilon_0\epsilon_0^2\int_{-\infty}^{\infty} dt\, u^2(t)\cos^2(\omega_L t)\int_{-\infty}^{\infty} dydz g^2(y,z) \\ &\approx \frac{c\varepsilon_0}{2}\epsilon_0^2\int_{-\infty}^{\infty} dt\, u^2(t)\int_{-\infty}^{\infty} dydz g^2(y,z) \\ &= \frac{c\varepsilon_0}{2}\epsilon_0^2 N_p\end{aligned}$$

The shape of the laser pulse is described by a 2D spatial gaussian and a 1D temporal gaussian with the characteristic parameters for laser beam waist $\omega_0$ and laser pulse duration $\tau_p$. The normalization term is thus:

(6) $g_{00}^2(x,y,z) \propto \exp\left(-\frac{2(y^2+z^2)}{\omega_0^2}\right)$

$$u^2(x,t) = \exp\left(-\frac{2\left(t-\frac{x}{c}\right)^2}{\tau_p^2}\right)$$

$$N_p = \frac{\omega_0^2\tau_p}{2}\sqrt{\frac{\pi^3}{2}}$$

For the purposes of brevity, we will solely address the magnitude of the phase shift. The phase can be further simplified as:

(7) $$\phi(\underline{r},t) \approx \frac{1}{\hbar}\frac{e^2}{\omega_L^2 m_e\gamma}\frac{E_L}{2c\varepsilon_0}\frac{\int dt' g^2(x,y,z)u^2(x,t')}{N_p} = \frac{1}{\hbar}\frac{e^2}{\omega_L^2 m_e\gamma}\frac{E_L}{2c\varepsilon_0}S(\underline{r},t)$$

$$= \frac{2\pi\alpha E_L}{\omega_L^2 m_e\gamma}S(\underline{r},t) = \frac{\alpha\lambda_L^2 E_L}{\gamma m_e c^2 2\pi}S(\underline{r},t) = \frac{\alpha\lambda_L^2 E_L}{2\pi E_e}S(\underline{r},t)$$

$$S(\underline{r},t)N_p = \int dt' \exp\left[-\frac{2}{\tau_p^2}\left(t'-\frac{x}{c}\right)^2\right]\exp\left[-\frac{2}{\omega_0^2}(z^2+y^2)\right]$$

By considering the positions $(x,y,z)$ as functions of time for a traveling electron, the phase shift is:

(8) $$\phi(\underline{r},t) \approx \frac{\alpha\lambda_L^2 E_L}{2\pi E_e}\frac{1}{N_p}\int dt' \exp\left[-\frac{2}{\tau_p^2}\left(t'-\frac{x}{c}\right)^2\right]\exp\left[-\frac{2}{\omega_0^2}(z^2+y^2)\right]$$

Phase is inherently difficult to measure. Instead, we measure the deflection caused by passing the electron through the laser beam. We expect the laser to roughly behave as a concave lens that is stronger in the axis perpendicular to the direction of laser travel. The momentum shifts are thus:

(9) $\delta\underline{p} = \hbar\nabla\phi(\underline{r},t)$

$$\underline{\nabla}\phi(\underline{r},t) = \frac{\hbar\alpha\lambda_L^2 E_L}{2\pi E_e}\frac{\nabla\int dt' g^2\left(x,y,z(t')\right)u^2\left(t'-\frac{x}{c}\right)}{N_p}$$

$$= \frac{\hbar\alpha\lambda_L^2 E_L}{2\pi E_e N_p}\int dt'\underline{\nabla}\left[g^2(x,y,z)u^2\left(t'-\frac{x}{c}\right)\right]$$

$$\Delta p_x = \frac{\hbar\alpha\lambda_L^2 E_L}{2\pi E_e N_p}\int dt'\left\{\frac{4\left(t-\frac{x}{c}\right)}{c\tau_p^2}\exp\left[-\frac{2\left(t-\frac{x}{c}\right)^2}{\tau_p^2}-\frac{2(y^2+z^2)}{\omega_0^2}\right]\right\}$$

$$\Delta p_y = \frac{\hbar\alpha\lambda_L^2 E_L}{2\pi E_e N_p}\int dt'\left\{-\frac{4y}{\omega_0^2}\exp\left[-\frac{2\left(t\prime-\frac{x}{c}\right)^2}{\tau_p^2}-\frac{2(y^2+z^2)}{\omega^2(x)}\right]\right\}$$

### 1.2 Assumptions

Application of the equations above make the following assumptions of the experiment:

1. The laser and electron beams interact in such a small region relative to the travel distance of the electron beam that the momentum and phase shifts are instantaneously applied to the

travel path of the electron beam. We have found this to be accurate in cases where the laser and electron pulse durations are at least shorter than 1 ns[2].

2. The gaussian beam profile has a best-fit to the fundamental transverse-electromagnetic ($TEM_{00}$) solution to the laser gaussian beam and ignores any fringes that may appear due to dust, mirror, or lens aberrations.

3. The photon oscillatory component (582 THz, ~1.7 fs period) truly plays no part on the timescales involved in the ponderomotive interaction (140 fs for the electrons to cross the interaction zone).

4. The effect can be averaged across varying electron pulse durations (due to fluctuations in electron counts) to be equivalent to an experiment performed at the average pulse duration.

### 1.3 Parameters Used

In Figure 1 of the main text, we show a simulation that provides a reasonable match to the data collected by experiment. The properties of the electron beam and laser beam used in that simulation are described in the table below.

**Table S1. Parameters for simulation from Figure 2 of the main text**

| Parameter | Used Value | Value Range |
|---|---|---|
| Magnification | 15kx | 10kx-20kx |
| Interaction to Sample | 7000 um | 7000 um[a] |
| Focus to Sample | -200 um | 0 to -630 um[b] |
| Interaction Size | 15 um | 12 to 24 um[c] |
| Laser Bunch | 212 fs | >212 fs[d] |
| Electron Bunch | 6200 fs | 5200 to 7200 fs[e] |
| Laser Focus | 12.5 um | 12.4 to 12.6 um[f] |
| Electron Focus | 1 nm | >1nm[g] |
| Electron Rotation | $-70^{o}$ | $-30^{o}$ to $-90^{o}$ [h] |
| Electron Astigmatism (⊥/‖) | 0.5 | 0.25 to 0.75[h] |
| Electron Energy | 30 keV | 30 keV |
| Laser Pulse Energy | 3.21 µJ/pulse | 3.21 µJ/pulse[i] |
| Laser Wavelength | 515 nm | 515 nm[d] |
| Laser Positional Offset (⊥) | 1.75 um | 1.75 um[e] |

[a]This is set by the position of the specimen stage
[b]This is the defocus range that the electron beam shifts by when moving from thermionic/Schottky mode to photoelectron mode
[c]As shown in Figure S4.
[d]Property of the laser
[e]As shown in Figure 2 of the main text
[f]As shown in Figure S2.
[g]Theoretical resolution limit of the SEM.
[h]As shown in Figure S4.
[i]Measured at a known position in the laser line

### Supplementary Note 2. Characterization of Experimental Parameters

#### Spatiotemporal Overlap

We have used previously established silicon photoexcitation experiments to synchronize the spatiotemporal overlap of electron and laser pulses[3]. A periscope with an adjustable lens mount is used to roughly align the laser into the electron beam field of view. This phenomenon is further explored in a forthcoming paper, though it is generally understood that photoexcitation induces transient electric fields that affect secondary electron generation[4]. These secondary and backscattered electrons are collected by an Everhart-Thornley Detector. The time zero experiments were performed on p-type low-B-doped silicon (Ted Pella p-type boron-doped Si <100> with 1-30 Ω·cm resistivity and 2 nm roughness) at 1 MHz, with the photoexcitation fluences set to 130 μJ/cm$^2$, well beyond the picosecond damage limit of silicon.

Shown below are a few of the experiments we have performed when attempting to locate this timepoint down to a precision of 0.1 mm (333 fs). We have primarily focused our attempts on varying the probe UV pulse energies, which should theoretically change the electron beam pulse duration and provide a pivot point which we can pin time zero to.

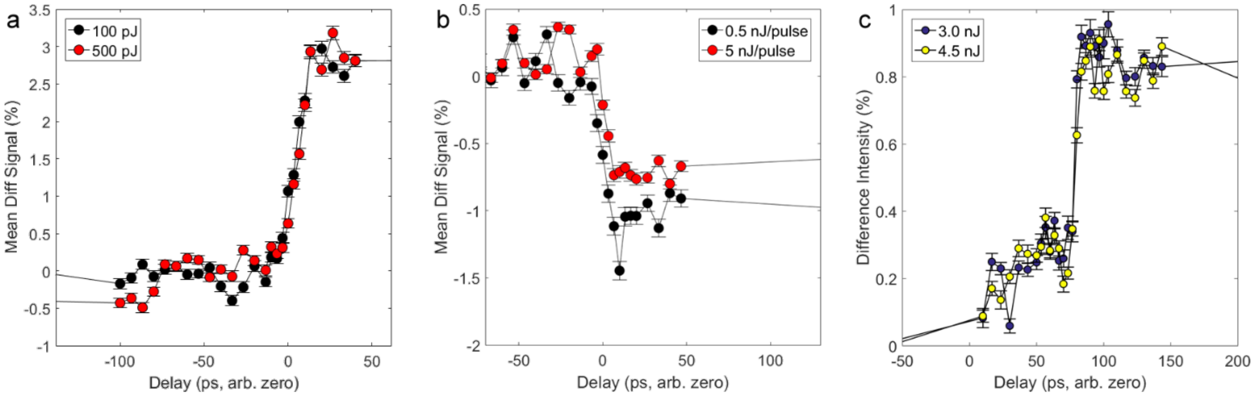


**Figure S1. Attempted probe UV pulse energy variation experiments at three different energy differences and in three different locations on intrinsic Silicon (lightly B-doped).** As can be observed from all three of these experiments, there is not a large enough difference in the photoexcitation responses of the underlying silicon to warrant any pinning of time zero. **a,** shows experiments performed between 100 pJ and 500 pJ UV pulse energies, **b,** shows the difference between 500 pJ and 5000 pJ, and **c,** the difference between 3000 pJ and 4500 pJ. Indeed, the second experiment (**b**) seems to suggest the opposite of theory

This suggests one of the three following theories:

1. The electron pulse is laser-limited out to 5 nJ/pulse. This places time zero at the middle of the response curve.
2. The phenomenon we are observing is material limited. This places time zero at the beginning of the response curve.
3. Some constraint in the electron microscope pins the electron pulse duration to a specific value which is highly insensitive to the number of electrons generated at the cathode. This places time zero at the middle of the response curve.

Given that other publications have clearly shown 1 and 3 to not be the case, we reasoned that time zero was at the beginning of the curves[3-5]. The ponderomotive experiments verified this hypothesis.

Laser Parameters

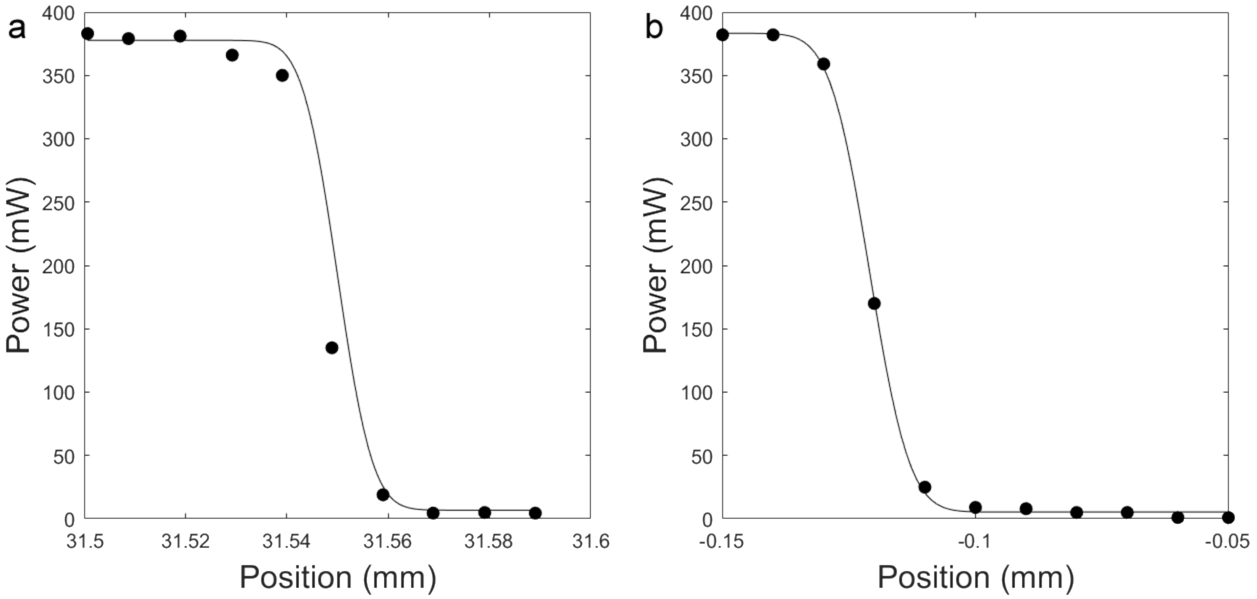


**Figure S2. Size of the pump laser beam in the two directions perpendicular to the Poynting vector. Beam sizes are determined with a silicon knife edge within the microscope. a,** The plot for the direction parallel to the electron beam. The size measured here is 12.6 microns. **b,** The plot for the direction perpendicular to both the Poynting vector of the laser and the electron beam. The size from the fit is 12.4 microns.

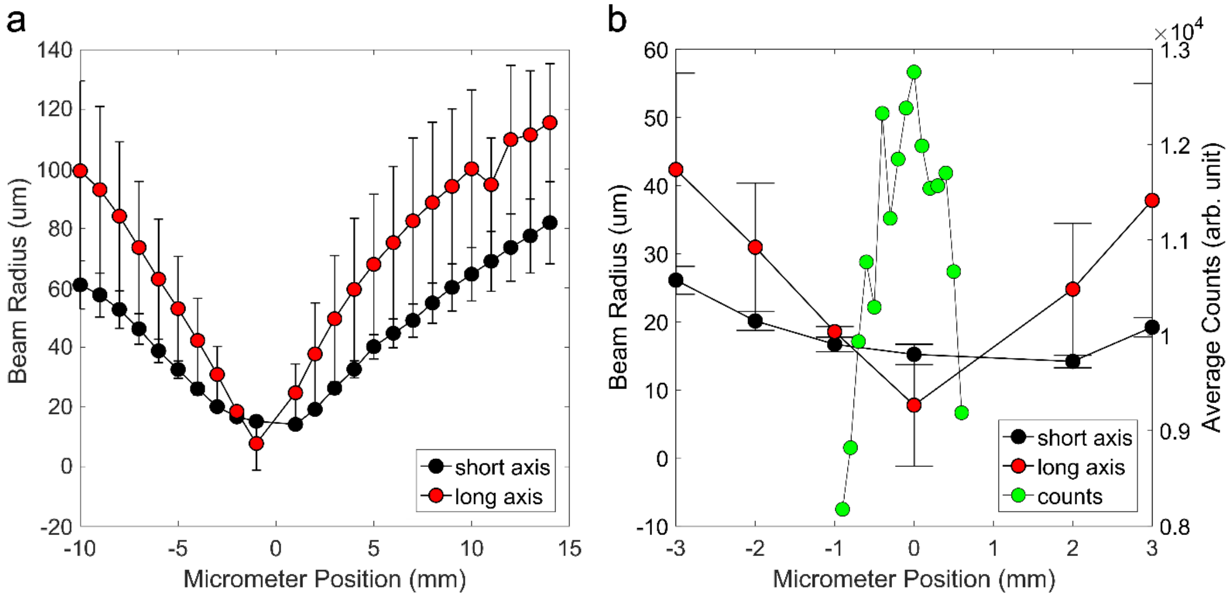


**Figure S3. Size of the UV laser beam striking the ZrO-tungsten cathode. a,** The size is split into two directions, with the long axis in the direction parallel to electron beam travel and the short axis perpendicular to the direction of electron beam travel. Barring vibrations and drift, the beam focus is generally placed on the cathode. Beam sizes are determined with an Ophir Optronics SP932 beam profiler. **b,** The counts sensitivity to the focal point position at a fixed brightness and contrast value on the detector. This provides a good description of the behavior of the electron beam, though the zero on the counts axis corresponds to the end of the dynamic range of the detector.

### Electron Beam Parameters

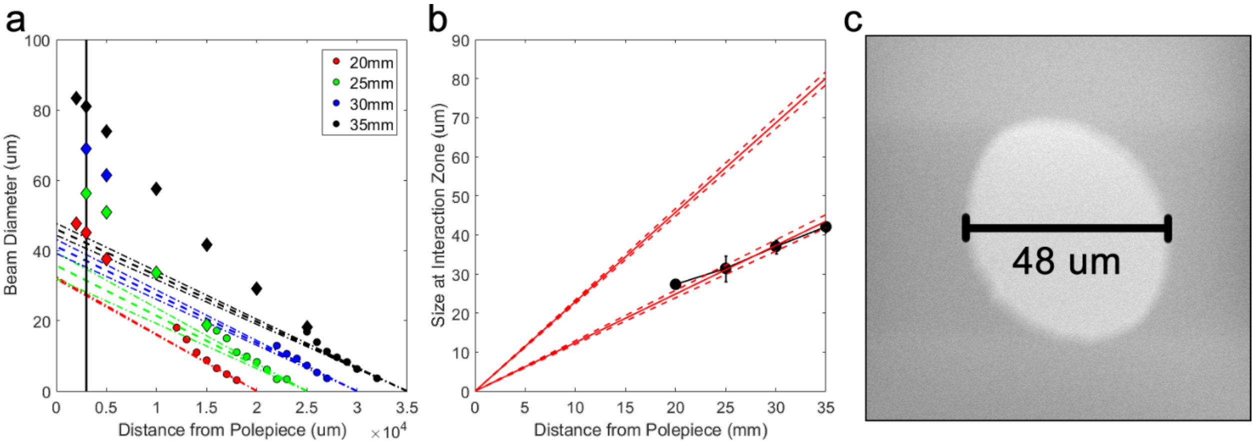


**Figure S4. Size of the electron beam at various distances from the focal point. a,** The electron beam ideally linearly increases in size as it moves further away from the working distance (focal point). The working distance is set to a specific position (indicated by color) and measurements of the beam size are acquired by imaging a small piece of dust and dividing the resulting size dimensions by 2. These dimensions are plotted as a function of distance from the polepiece. As can be seen, there is a nonlinearity in the data. **b,** When the 1000 micron aperture is used, the size of the electron beam in the interaction region (7 mm) is 12 to 24 microns. These two values are derived from the information found by the non-linear data split into a far-field set and a near-field set. The resulting beam size must be somewhere in the middle. This value serves as a useful guide for simulations work. **c,** A description of the electron beam size 3mm from the polepiece. The true size of the electron beam must be divided by two. The electron beam asymmetry is due to electron beam not being constrained by any objective aperture and a misalignment in the column discovered after the experiment.

### Secondary Control

To ensure the effect we observed was central to the interaction region between the laser and the electron beam, we performed one additional control. The specimen was moved 23.5 mm away from the interaction zone. If the observed effect was dependent on the surface of the specimen, then the $t_0$ position would move. Here, we observed no difference in $t_0$, ensuring that the effect we observed is localized to the laser-electron interaction zone.

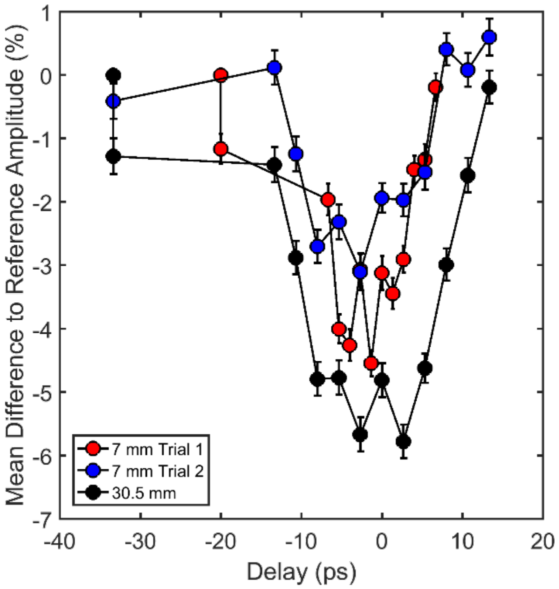


**Figure S5. Interaction control to ensure no surface effect.** The same trials from the 0.5 nJ/pulse experiments from Figure 4 of the main text are repeated here, including the reference images. A third experiment involving the moved specimen set to 30.5 mm away from the polepiece is also overlayed on the image. We can see that $t_0$ has not moved by an expected 23.5 mm on the delay stage (more than 100 ps).
